# Distributed hierarchical filters partition the Danionella vocal repertoire

**DOI:** 10.64898/2026.03.04.709502

**Authors:** Jörg Henninger, Maximilian Hoffmann, Mykola Kadobianskyi, Johannes Veith, Caroline Berlage, Antonia Groneberg, Daniil Markov, Lisanne Schulze, Ana Svanidze, Leonard Maler, Benjamin Judkewitz

**Author notes:** these authors contributed equally to this study (Jörg Henninger, Maximilian Hoffmann).

## Abstract

Vocal repertoires consist of a limited number of call types, even though their sounds can vary continuously in form. Where in the brain receivers partition this variation into discrete types is contested: different accounts place the emergence of categories in the cortex/pallium or already in the midbrain. Because recordings have been limited to isolated regions, it remains unknown whether this partitioning is confined to a single region at all, occurs in parallel across many, or is built up along the pathway. Here we present the first whole-brain, cellular-resolution analysis of how conspecific vocalizations are categorized in a vertebrate, using the glassfish *Danionella cerebrum*. We find that categorization is neither pallial nor localized. It begins at the first central auditory nuclei, much earlier than previously thought, where a continuous morph between social and non-social sounds evokes an abrupt switch in population activity. The midbrain then sharpens these representations, and the thalamus filters for species-typical pulse rates while splitting call duration into two discrete populations. Only downstream of this shared code do the sexes diverge in forebrain social nuclei. Call type partitioning is thus distributed and hierarchically assembled, rather than read out at any one stage.

## Introduction

Acoustic communication is a prime example of how nervous systems partition a continuous signal space into discrete, behaviorally meaningful types. Receivers as different as insects, frogs, birds and mammals can sort calls into distinct kinds that lead to different responses, even where the types grade into one another acoustically ^1–3^. Where in the auditory pathway this partitioning arises has been contested. Categorical perception is classically placed in the cortex, the endpoint of a pathway in which subcortical stations extract acoustic features and pass them upward for integration ^4,5^. Yet neurons selective for call types have also been found closer to the periphery: in the anuran midbrain ^6^ and the bat inferior colliculus ^7^. It therefore remains unclear whether classification happens at a single stage, is repeated redundantly across many, or is built up hierarchically. This question cannot be settled without observing the whole pathway.

However, despite substantial progress in select vertebrate models, recordings and manipulations have largely been confined to a subset of brain regions, yielding detailed but severely undersampled accounts of social signal processing. Brain-wide approaches based on immediate early gene expression offer unbiased and broad coverage ^8^, but their slow, integrative nature precludes stimulus-resolved comparisons and can conflate stimulus-evoked activity with state-, context-, and motor-related activity. Calcium imaging in larval zebrafish has mapped auditory responses to synthetic sounds at cellular resolution ^9^, but zebrafish do not produce social vocalizations and the larval brain lacks the mature circuits that link auditory processing to social behavior. As a result, we still lack brain-wide recordings revealing how socially relevant sounds are represented and transformed across the brain. It also remains unclear where along the hierarchy selectivity for conspecific vocalizations first becomes segregated from non-social, ambient sounds, and at what stages sex-dependent differences in call representation emerge – whether from peripheral filtering ^10^ or from divergent downstream computations of valence in social decision-making circuits.

To close this gap, we exploit the distinctive features of *Danionella cerebrum*, a vocalizing glassfish uniquely suited for such comprehensive, brain-wide investigations (Fig. 1a) ^11–13^. Its small, optically accessible brain enables non-invasive, volumetric imaging of neuronal activity across all brain regions simultaneously (Fig. 1b) ^13–17^. Male *Danionella* vocalize during agonistic encounters and courtship by drumming their swim bladder to emit temporally structured pulse bursts generated at ∼60 or ∼120 Hz (Fig. 1c) ^11,18,19^. Bursts vary in duration, occurring either as short bursts (typically 50–150 ms, comprising 6–20 pulses) arranged in sequential series, or as single prolonged bursts lasting from seconds to several minutes (Fig. 1c). These vocalization patterns provide an experimental handle for studying auditory processing of ethologically relevant sounds. Together, these properties make *Danionella* an ideal system for studying how brain-wide circuits detect social sound and categorize call types. By recording across the entire brain, we find that call types are not read out at any single stage, but assembled along the pathway: separated from other sounds already at the first central auditory nuclei, filtered and split by the thalamus, and only then valued differently by the two sexes.

**Figure 1.**
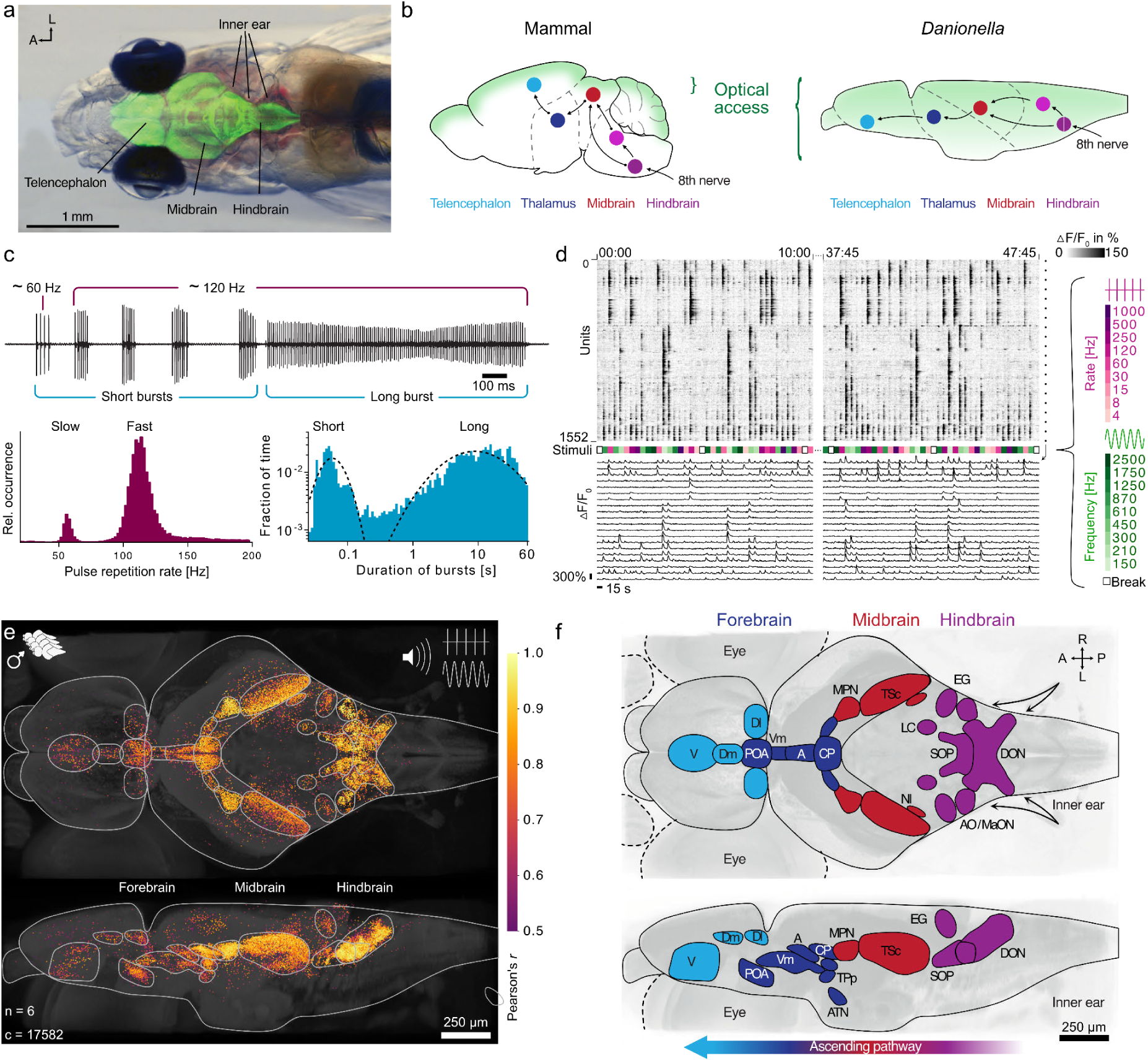
Whole-brain auditory responses in *Danionella cerebrum*: **a**, Photograph of an adult transgenic pigmentation-knockout male. Overlay in green: *in vivo* laser scanning reflectance contrast section of the brain, serving as anatomical reference. **b**, Schematic of a typical mammalian and a *D. cerebrum* brain. Transparent *D. cerebrum* offers noninvasive optical access to all major auditory processing centers. **c**, Top: Trace of acoustic vocalizations produced exclusively by males – temporally patterned pulse bursts at two repetition rates. Bottom left: Histogram of vocalization pulse repetition rates, with two prominent peaks at ∼60 and ∼120 Hz. Bottom right: Histogram showing the fraction of time contributed by bursts in each duration bin. The dashed line is a maximum-likelihood fit of a two-component log-normal mixture with maxima at 0.06 and 8.9 seconds. **d**, Temporal dynamics of auditory responses. Top: Rastermap of neuronal responses to pulse bursts and pure tone stimuli ordered by response similarity. Center: Stimulus timing and properties (see legend at right). Bottom: Example calcium traces (ΔF/F) from equally-spaced rastermap locations (black dots, right margin). Left: First 10 minutes of recording; right: final 10 minutes. **e**, Orthogonal projections of whole-brain auditory responses aggregated across 6 male fish. Each dot represents a responsive cell projected onto the male reference template. Color indicates the degree of entrainment with the acoustic stimulation, measured by Pearson correlation coefficient *r* between the stimulus regressor and the neuronal activity. **f**, Schematic of major acoustically responsive regions. Abbreviations: A, anterior *thalamic nucleus;* AO/MaON, *anterior octaval nucleus / magnocellular octaval nucleus*; ATN, *anterior tuberal nucleus*; CP, *central posterior thalamic nucleus*; Dm, medial zone of the dorsal telencephalon; Dl, lateral zone of the dorsal telencephalon; DON, *descending octaval nucleus*; EG, granular eminence of the cerebellum; LC, *locus coeruleus*; MPN, *medial pretoral nucleus*; NI, *isthmic nucleus*; POA, *parvocellular preoptic nucleus*; SOP, *secondary octaval population*; TSc, *Torus semicircularis*; TPp, periventricular nucleus of the posterior tuberculum; V, ventral area of the subpallium; Vm, ventromedial thalamus; one uncharted diencephalic region ventrolateral of CP.

## Results

### Brain-wide map of responses to social and non-social sound

To obtain a brain-wide map of responses to social and non-social acoustic cues, we performed volumetric calcium imaging of transgenic male *Danionella cerebrum* expressing nuclear-localized GCaMP6s across the brain ^14^. We took advantage of the simple parametric structure of their vocalizations – pulses repeated at stereotyped rates with varying burst duration – that enables a systematic exploration of the stimulus space. As a first step, we sought to identify acoustically responsive neurons across the brain. To elicit responses in as many auditory neurons as possible across the auditory pathway, we designed a broad stimulus set. This included pulse sequences (bursts) mimicking *D. cerebrum* vocalizations at various repetition rates, as well as pure tones at different frequencies, which have traditionally been used as elementary stimuli in auditory research ^20^. The brain-wide response maps revealed a broad sensitivity to pulse trains from 4 Hz up to ∼500 Hz repetition rate and to pure tones from 150 Hz up to ∼500 Hz (Fig. 1d; Fig. S1,2). To compare the spatial distribution of auditory responses across individuals, we registered the recordings to our reference brain atlas (^21^; Fig. 1e). The resulting aggregate map showed dense clusters of acoustically responsive neurons distributed across the entire brain. Mapping the functional imaging data onto canonical brain regions, we identified 19 acoustically responsive brain areas distributed across all major divisions of the brain (5 hindbrain, 1 cerebellar, 3 midbrain, 7 diencephalic, 3 telencephalic nuclei). These regions span the canonical ascending auditory pathway, from primary sensory processing in the hindbrain through midbrain and thalamic relay nuclei to higher-order integration in the telencephalon, and define a set of candidate nodes through which social sounds can influence behavior (Fig. 1f).

### Neuronal responses to social and non-social sound segregate early

We next asked where along the auditory pathway responses to vocalization-like pulsed sounds become segregated from responses to non-social pure tones. To analyze this, we plotted the trial-averaged responses (calcium signal ΔF/F) for all units across stimuli and sorted cells by the center-of-mass of their tuning curves: most cells responded selectively to either pulse trains or pure tones, with only a small fraction showing strong responses to both stimulus types (Fig. 2a). To quantify this selectivity, we defined a pulse/tone preference index ranging from -1 (exclusive pulse-train selectivity) to +1 (exclusive tone selectivity; see Methods). The distribution of this index showed one peak at each of the extremes: 21.0 ± 2.6 % (mean ± SD) of units (i < -0.5) preferred pulse trains and 54.4 ± 14.4 % (i > 0.5) preferred tones. The remaining 24.6 ± 14.5 % (−0.5 ≤ i ≤ 0.5) displayed mixed selectivity (Fig. 2b).

**Figure 2.**
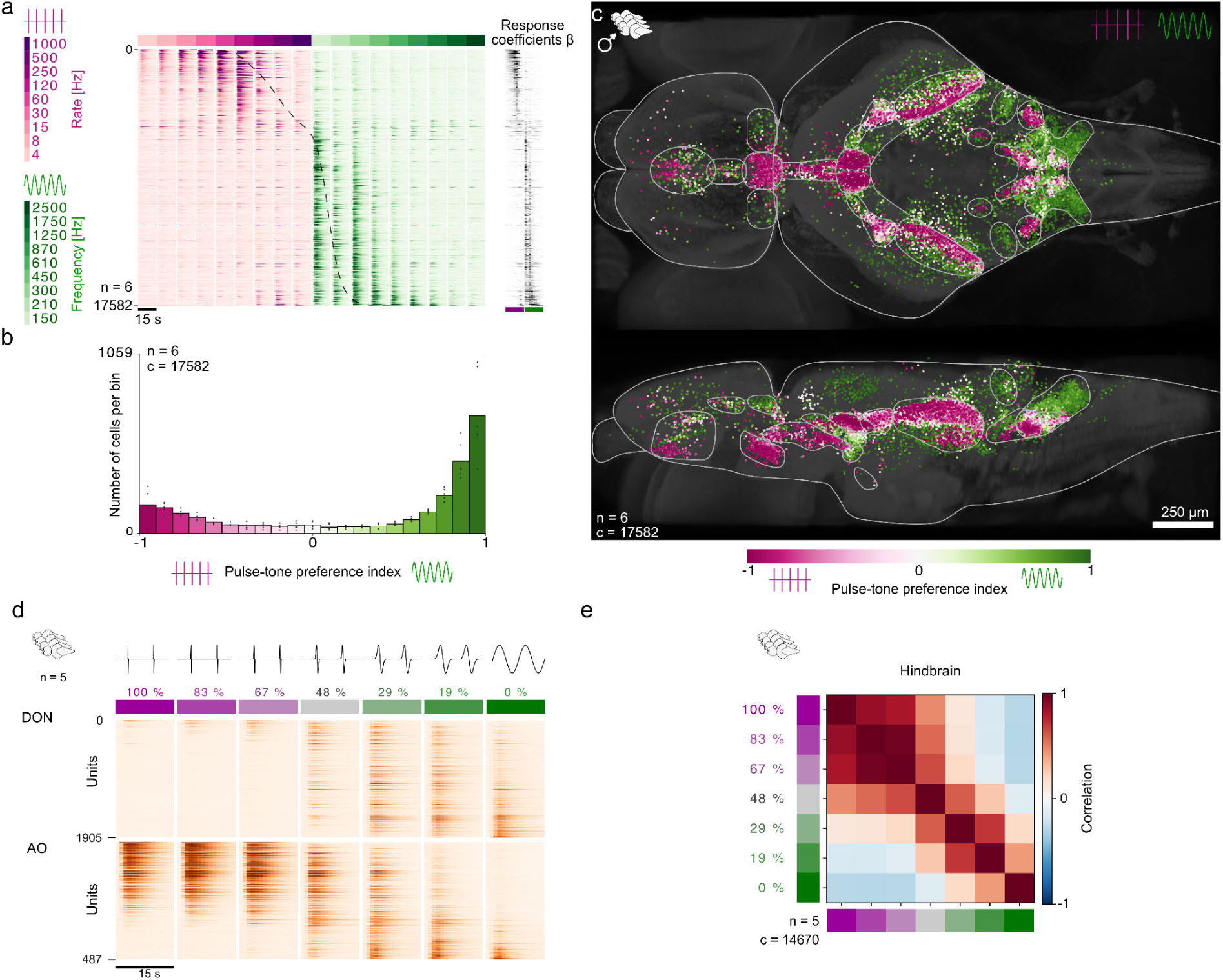
Tonal and vocalization-like sounds are segregated already at the first hindbrain nodes: **a**, Population response profiles to pulse bursts (magenta) and pure tones (green). The heatmap shows trial-averaged calcium responses (ΔF/F) for individual neurons (rows) across pulse bursts and tones at various frequencies (columns), pooled across 6 fish and sorted by the center-of-mass of individual tuning curves to reveal pulse/tone preference gradients. Stimulus composition shown above with legend at left. Right: Regression weights corresponding to the neuronal response traces at left. **b**, Distribution of pulse/tone preference indices across the population. Histogram bins show mean response counts (boxes) with individual fish contributions overlaid (black dots; n=6 fish). Values near -1 indicate dominant pulse preference; values near +1 indicate dominant tone preference. **c**, Spatial distribution of pulse-preferring (magenta) and tone-preferring (green) neurons aggregated across 6 fish and color-coded by pulse-tone preference index. **d**, Hindbrain population response profile to the pulse-tone morph. The heatmaps show trial-averaged calcium responses (ΔF/F) for individual neurons from the two primary hindbrain nuclei, the descending octaval nucleus (DON) and the anterior octavolateral nucleus (AO), pooled across 5 male fish and sorted by the center-of-mass of individual tuning curves. **e**, Hindbrain stimulus correlation computed from responses to the pulse-tone morph and averaged across 5 male fish.

Auditory systems are usually thought to transform neuronal response patterns from dense representations in primary afferents to feature-selective sparse representations in midbrain regions ^6,22–26^. We therefore expected that selectivity for pulses and tones would first appear in the midbrain torus semicircularis, the teleost equivalent of the mammalian inferior colliculus ^27^. Instead, discrete and spatially segregated pulse- and tone-evoked responses were already evident in first-order hindbrain nuclei receiving auditory afferents, including the descending and magnocellular octaval nuclei, and the secondary octaval population (DON, AO/MaON, SOP; Fig. 2c; Fig. S3). This segregated tuning emerged unexpectedly early and extended across much of the brain. Many auditory regions responded to both pulses and tones, yet with distinct topographic organization within each region (Fig. S3; S4). We noted prominent mixed-selectivity clusters in the hindbrain secondary octaval population (SOP), and in the midbrain torus semicircularis (TSc) and medial pretoral nucleus (MPN), and less dense clusters in the thalamic nuclei and the preoptic area (Fig. S3).

To determine whether the transition between response patterns to pulses and tones occurs gradually or categorically, we designed a stimulus that morphs from a pulse-burst with 120 Hz repetition rate to a 120 Hz pure tone (Fig. 2d). Across this continuum, neuronal responses did not change gradually but switched sharply at intermediate stages (Fig. 2e; 29-48% morph, see Methods).

In summary, tones and vocalization-like pulse bursts engage segregated neuronal populations from the earliest hindbrain processing stages onward.

### Thalamus gates calls with species-specific pulse-rates

Having established that pulse-tone segregation arises early in central auditory processing, we next asked how the auditory system encodes the specific acoustic features that define conspecific vocalizations. *D. cerebrum* calls are characterized by two species-specific pulse repetition rates (∼60 Hz and, predominantly, ∼120 Hz) and stereotyped temporal patterning (Fig. 1c) ^11^. This structure suggests two candidate cues for conspecific detection and call type discrimination: pulse repetition rate and burst duration. To test this idea, we first focused on the pulse repetition rate by varying pulse rate from 15 up to 250 Hz and analyzing the response profiles across brain regions.

Trial-averaged response heatmaps and their corresponding covariance maps revealed that auditory hindbrain neurons are broadly sensitive to pulse rate, with the exception of a few cells narrowly selective to < 15 Hz and > 250 Hz (Fig. 3a,b; S5). Midbrain TSc neurons were more selective than hindbrain neurons (Fig. 3a,b; Fig. S5), consistent with previous reports ^6,24–26^. The population remained sensitive across the full range of tested rates, with overrepresentation of 105–120 Hz that matches the prominent mode of the natural distribution of vocalization rates (Fig. S5; 1c). For each neuron, we characterized the best rate as the pulse rate eliciting the maximum response, and mapped these values across recording sites to assess topographic organization. Both hindbrain and midbrain areas exhibited mild topographic best-rate organization along the anterior-posterior axis (Fig. 3d). In contrast to the full representation of all rates tested in the hind- and midbrain, the first thalamic nucleus, the *central posterior thalamic nucleus* (CP th.), was almost exclusively peak-tuned to 105–120 Hz pulse rate with comparable selectivity to the midbrain (Fig. 3a-d; S5). This specific selectivity for vocalization-typical rates was maintained in most downstream regions, particularly in the subpallial ventral zone of the ventral telencephalon (Vv; Fig. S4), and was robust across all animals (n=9; Fig. 3a-c; S6; with the exception of a cluster of > 200 Hz selective cells in the rostral ventromedial thalamus (VM th.) and the subpallium). Interestingly, neurons in the preoptic area (POA) – a central node of the conserved social behavior network known to be associated with aggressive, affiliative, and sexual behavior ^28–30^ – were more coarsely tuned to vocalization rate than CP thalamic neurons, suggesting that the POA receives auditory input from additional sources beyond CP thalamus (Fig. 3c; S5). Many neurons in TSc, CP thalamus, VM thalamus, POA and Vv peak-tuned to 120 Hz also displayed weak secondary peaks at 60 Hz, which is the less prominent vocalization rate (Fig. 1c; Fig. S5). In sum, while midbrain neurons still tile a broad range of pulse rates, the thalamic CP acts as a narrow filter that encodes only signals with species-typical vocalization rates.

**Figure 3.**
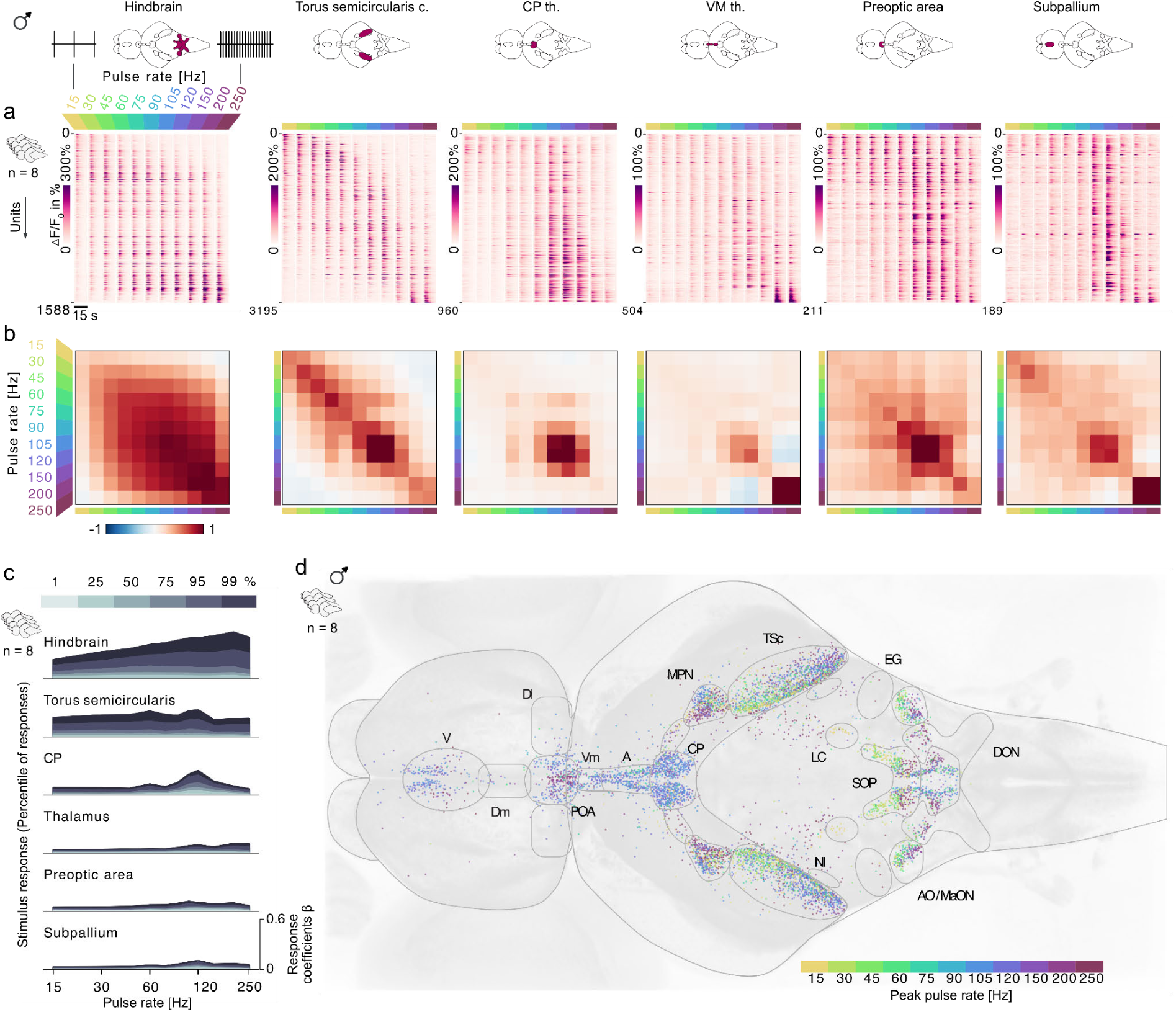
Thalamus gates calls with species-specific pulse rates: **a**, Population-level regional response profiles, aggregated across 8 fish. Heatmaps show trial-averaged calcium responses (ΔF/F) for individual neurons (rows) across pulse repetition rates from 15 to 250 Hz (columns) in six auditory brain regions (labeled above each panel). Neurons are sorted by center-of-mass of individual response curves to reveal rate selectivity gradients. **b**, Normalized stimulus covariance maps computed from regional neuronal responses and averaged across fish. Normalized to the 99.7% percentile. **c**, Regional response magnitude distributions. Percentile curves (1st–99th percentile) of stimulus-evoked response amplitudes by brain region, showing differential sensitivity across auditory regions. Y-axis shows response coefficients 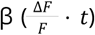. **d**, Spatial distribution of repetition rate preferences. Top view projection showing anatomical locations of auditory neurons color-coded by their best-response to pulse repetition rate, revealing topographic organization of rate selectivity.

### Thalamus partitions call duration

After pulse rate, the second prominent feature of *D. cerebrum* vocalizations is burst duration. We therefore examined whether short and long continuous bursts elicit different responses. Again we recorded brain-wide neuronal responses, this time focusing on bursts at 120 Hz pulse rate with durations ranging from 8 ms up to 4s (corresponding to 2 to 480 pulses). In the hindbrain, trial-averaged responses increased uniformly with burst duration (Fig. 4a–d; S9) ^31–33^. The midbrain showed response profiles similar to the hindbrain, but we noted a small population in the posterior TSc specifically tuned to short bursts (Fig. S10). Remarkably, the vocalization-rate selective thalamic CP contained two spatially distinct, equally sized populations with peak selectivity to short and long bursts, respectively. Similar but less narrow short- and long-burst selectivity was found in the thalamic VM and the POA, while the anterior thalamus (A) exhibited short-burst selectivity and the hypothalamic anterior tuberal nucleus (ATN) displayed predominantly weak long-burst selectivity (Fig. S10). Vv was exclusively peak-tuned to long-duration bursts. Similarly to CP, the parvocellular POA displayed weaker but defined duration-selective topography, with short-burst-selective cells concentrated in the anterior and long-burst-selective cells in the posterior part.

**Figure 4.**
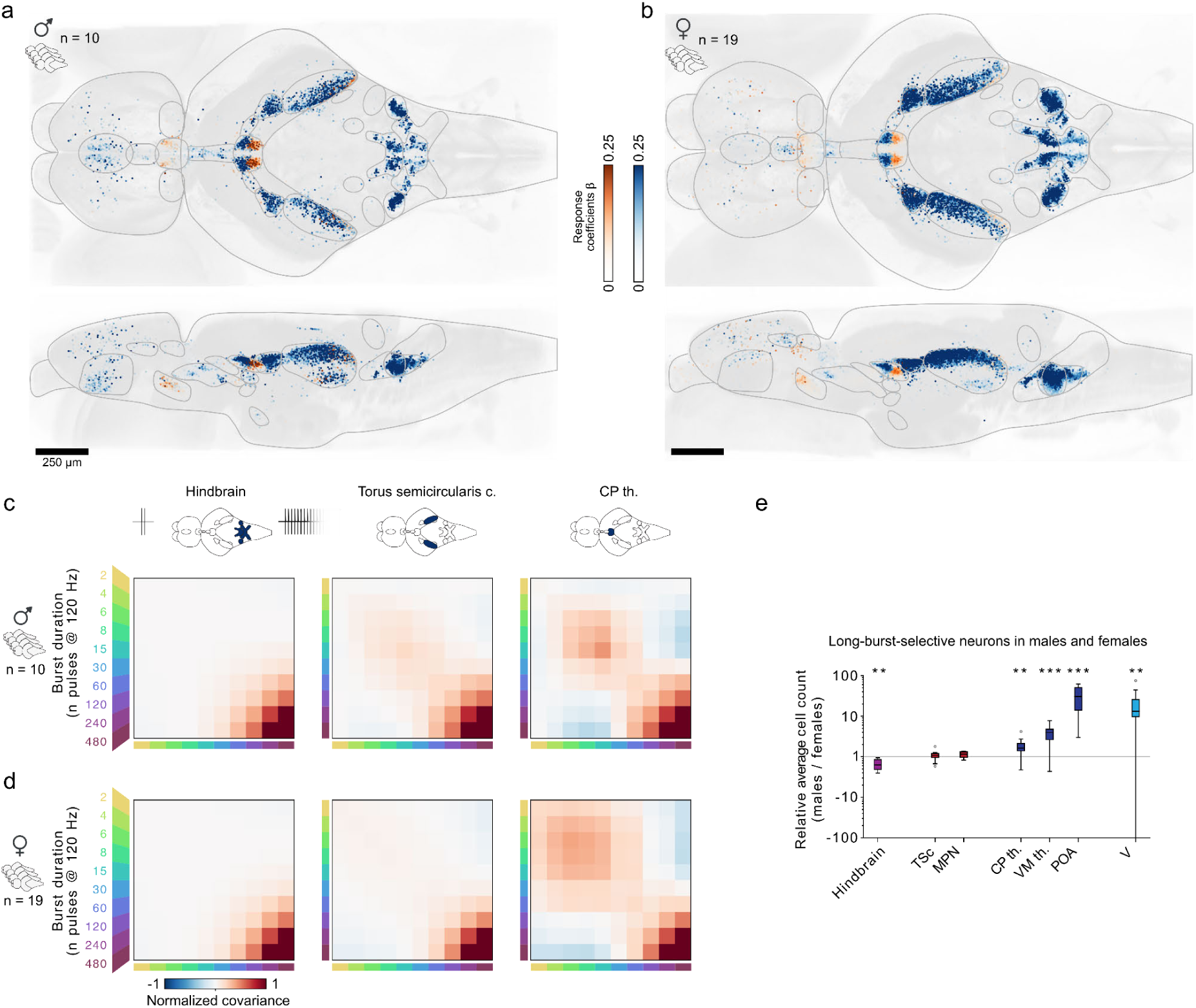
Thalamus partitions call duration and forebrain activity is sexually dimorphic: **a**, Male response magnitudes to short (orange: 2-30 pulses) and long bursts (blue: 120-480 pulses), pooled across recordings (n=10). **b**, Female response magnitudes to short and long bursts (n=19). **c**, Male stimulus covariance maps computed from regional neuronal responses to pulse bursts varying from 2 to 480 pulses per burst and averaged across 10 fish, normalized to the 99.7th percentile. **d**, Female normalized stimulus covariance maps, averaged across 19 fish. **e**, Relative numbers of long-burst-selective cells in males and females by region (male counts divided by female median count). Regional counts were compared between sexes using a Mann-Whithney U test; significant differences were observed in the hindbrain, CP th., VM th., POA, and V (hindbrain: p=0.0016, CP th.: 0.0016, VM th.: 0.0009, POA: 0.0002, and V: 0.0042, respectively).

Whereas hindbrain responses scale continuously with burst duration, CP splits this continuum into two discrete populations, one for short and one for long bursts. Where earlier stages separate social from non-social sounds (Fig. 2e), CP is the first to partition conspecific calls into distinct call duration types (Fig. 4c,d).

### Sex-specific sensory representations emerge in the forebrain

Only males produce social sounds, but both sexes are receivers. Because identical acoustic features can be associated with different behavioral outcomes in females and males (for example, courtship versus aggression), those features could plausibly carry sex-dependent valence ^34,35^. Sex differences could enter at either end of this pathway. If they originate peripherally ^10^, males and females should differ already in the early feature code. Alternatively, if the sexes share a common sensory representation and differ only in how they value it, early tuning should be sex-invariant and divergence should appear only in forebrain circuits that couple sensory categories to social action.

To distinguish these possibilities, we compared how males and females encode pulse rate and burst duration across the auditory pathway. Rate response profiles were qualitatively similar in the male and female hind- and midbrain, except that more neurons were responsive in the hindbrain of females overall. However, marked sex differences emerged in downstream forebrain regions: in females far fewer neurons responded to long 120 Hz bursts in thalamic nuclei (CP, VM), the POA, and the subpallial Vv. Forebrain response magnitudes to 120 Hz bursts were also weaker overall (Fig. S5–8).

In line with the rate selectivity, duration response patterns were qualitatively similar in the hindbrain, in the midbrain, and CP thalamus, but diverged in the forebrain (Fig. 4; S9; S10). The proportion of short-burst tuned neurons remained similar across sexes in all forebrain regions, while we observed a broadening of short burst-selectivity in the male forebrain (Fig. S9–11). However, markedly more neurons responded to long bursts in males than in female forebrains: CP th.: 1.6x (*p*=0.002); VM th.: 4x (*p*=0.001); POA: 30.5x (*p*=0.0002); V: 13.3x (*p*=0.004) (Fig. 4a-b; 4e; S11).

Together, these results show that sexual dimorphism in burst-duration selectivity emerges downstream of the midbrain: substantially fewer neurons responded to long bursts in the female thalamic, preoptic, and subpallial regions, suggesting that identical acoustic features acquire sex-specific valence in forebrain social circuits.

## Discussion

Here we provide the first unbiased, cellular-resolution analysis of social acoustic processing across a whole adult vertebrate brain. Sorting communication sounds into behaviorally meaningful types is not accomplished at any single stage, but built up progressively along the auditory pathway. Vocalization-like pulse trains are set apart from other sounds already in the brainstem; the midbrain sharpens this selectivity and extracts temporal features; the thalamic CP filters for the species-typical pulse rate and splits burst duration into discrete populations; and only in the forebrain do these shared sensory categories acquire sex-specific responses. Call type categorization is thus a distributed, hierarchical process rather than the property of one region.

### Segregation of pulsed and tonal sounds in the Hindbrain

Our whole-brain mapping reveals that social signal segregation begins far earlier than previously shown ^6,24–26^. While classical models acknowledge the role of the hindbrain in transforming dense peripheral representations into sparse, feature-selective rate codes, they typically assign complex scene analysis – such as distinguishing calls from other environmental sounds – to midbrain or forebrain circuits ^4,26^. Instead, we find that *D. cerebrum* separates vocalization-like pulsed sounds from tonal sounds at the very first hindbrain nuclei.

### Selectivity for vocalization rate and duration in the central posterior thalamus

The auditory midbrain and thalamus play distinct but complementary roles in vocal processing. While the midbrain TSc acts as a broadband feature processor responding to a wide range of pulse rates and tone frequencies, the thalamic CP functions as a gatekeeper for social sounds: it responds selectively to conspecific vocalization rates. This represents a pronounced sharpening from midbrain to thalamus, whereby the CP selectively transmits only species-specific signals to downstream forebrain networks. Crucially, the CP adds another layer of selectivity by transforming vocalization duration into two discrete categories. While hindbrain responses scale continuously with stimulus length, the CP contains two spatially segregated populations – one tuned to short bursts, the other to long bursts.

Anatomically, the CP is well-positioned to transmit conspecific signals to multiple social brain structures. In other teleosts, it receives direct input from the midbrain TSc, responds to acoustic stimuli, and projects to structures known to be involved in social behaviors, including the ATN, the POA, and Vv ^36–39^ – components of what has been proposed as a conserved social decision-making network ^28,40,41^. This positions the CP as a thalamic “social gate” that couples vocalization categories to conserved limbic circuits controlling agonistic and reproductive behaviors ^42,43^. The strong CP connectivity with the hypothalamus, preoptic area and subpallium ^38,39^ as well as the pallial amygdala (Dm) ^44,45^ identifies it as similar to components of the mammalian intralaminar thalamic nuclei ^46^. The direct projection from CP to Dm explains the observed pallial activity despite the lack of responses in the preglomerular complex, the dominant diencephalic relay to the teleost pallium ^47^. Vv contains components homologous to the mammalian septum and pallidum (^48^; Fig. S4). Comparable, but not homologous, architectures exist in mammals and birds, where the mammalian dorsal thalamic medial geniculate and the avian ovoidalis nuclei show selectivity for behaviorally relevant spectrotemporal patterns of social vocalizations, respectively (e.g., ^49,50^), suggesting that thalamic gating of communication signals is a computational feature across vertebrates. This aligns with emerging evidence that the teleost posterior thalamus serves as a multisensory integration hub where social cues across modalities converge (visual, auditory, mechanosensory) – a function speculated to be conserved across vertebrates ^41,51,52^.

### Sexually dimorphic neuronal responses in the forebrain

Sexually dimorphic responses to vocalizations do not arise in the hindbrain, midbrain, or CP, where response profiles are highly similar between sexes, but instead emerge selectively in distinct regions of the diencephalon (VM, POA) and subpallium (Vv). In these regions female responses to long bursts are markedly reduced compared to males. In vertebrates from fish to mammals, the preoptic area and its subpallial targets constitute a conserved circuit for the control of vocal production and sexually dimorphic motor behavior ^53–56^.

## Conclusion

By exploiting the optical transparency of *Danionella cerebrum*, we provide a brain-wide, cellular-resolution account of how social sounds are sorted into call types. The organisation we describe, from early separation of communication sounds through progressive refinement to a thalamic gate and sex-specific forebrain readout, maps onto conserved elements of the vertebrate social brain, suggesting that distributed, hierarchical categorization may be a general strategy for social communication across vertebrates.

## Methods

### Animal care and transgenic Danionella cerebrum

All animal experiments conformed to Berlin state, German federal and European Union animal welfare regulations and were approved by the LAGeSo, the Berlin authority for animal experiments. *D. cerebrum* were kept in commercial zebrafish aquaria (Tecniplast) with the following water parameters: pH 7.3, conductivity 350 µS/cm, temperature 26 °C. We used male and female adult fish between 4 and 17 months of age. All imaging data was acquired using transgenic fish expressing the nuclear-localised fluorescent calcium indicator GCaMP6s under the pan-neuronal HuC promoter.

### Whole-brain imaging

#### Optical setup

For whole brain imaging we employed a custom-designed volumetric oblique-plane microscope as described in ^14^, with the following modifications: we used a transmissive diffractive grating (Coherent LightSmyth T-1600-1060s-12.3×12.3-94) as diffractive element in the intermediate image plane and updated the camera (Teledyne Kinetix). Effectively, this imaging system enabled a resolution of 2.3 ± 0.7 × 2.2 ± 0.91 × 9.4 ± 3.6 μm^3^ (FWHM, n = 9911 beads) at 1 Hz volume rate across a FOV of 2.2 × 0.8 × 0.9 mm^3^, along x, y, and z, respectively.

#### Image postprocessing

Analysis of neural volumetric time-lapse recordings requires pre-processing as detailed in ^14^. In short, we correct each camera frame for the static camera background. Because our microscope natively acquires each volume in a sheared coordinate system, each volume was de-sheared and deconvolved by 10 iterations of Richardson-Lucy deconvolution using a kernel empirically estimated from a fluorescent bead sample. Finally, all volumes were motion-corrected using affine and non-rigid registration to a template volume. Code for these preprocessing steps is available at: github.com/danionella/opm_unshear and github.com/danionella/warpfield.

#### Whole-brain imaging under acoustic stimulation

To ensure faithful playback of stimuli, the imaging setup was sound-calibrated once a day prior to the experiments (see “Calibrated acoustic stimulation” section). Fish were anesthetized in 120 mg l^−1^ buffered MS-222 and subsequently placed on a preformed, size-matched agarose mould, which allowed the gill covers to move freely, and immobilized with 2% low-melting-point agarose (melting point 35 °C). A flow of fresh, aerated aquarium water was delivered to their mouth through a glass capillary. Water-temperature was maintained at 25–26 °C during the experiment by a PID-controlled heating system consisting of a small thermometer placed in the mould below the fish and a heating element at the bottom of the tank.

After experiment onset, the intensity of the excitation beam was gradually increased over 4 minutes up to a final power of ∼3 mW (below the objective) to allow for slow habituation. Recording and read-out of one plane took 2.2 ms. We imaged 450 planes spanning 900 μm at 1 Hz volume rate. After post-processing and unshearing (which involves upsampling along Y and binning along Z) the datasets consisted of volumes with a size of, typically, 2650 × 1388 × 186 px (XYZ).

### Calibrated acoustic stimulation

Acoustic small tank experiments are complicated by the fact that tank geometry and receiver position both affect the acoustic signals experienced due to reverberations at the tank walls and the resulting superposition of reflected acoustic waveforms. We addressed this by recording each speaker’s impulse response at the fish’s location in terms of sound pressure and particle acceleration, effectively enabling precise control of these acoustic components experienced by the fish during stimulation ^57^.

#### Pressure and acceleration measurements

We characterized the acoustic playback properties of our experimental tank using three hydrophones that were positioned at the head position of the imaged fish (one centered and one to each side), aligned with the two opposing playback speakers. (Aquarian Scientific AS-1, preamplifier: Aquarian Scientific PA-4, acquisition: NI-9231 sound and vibration module, National Instruments). Pressure was measured with the hydrophone centered at the head position of the imaged fish. Particle acceleration was measured indirectly by the pressure gradient between two hydrophones positioned left and right of the fish head position. The distance between the hydrophones was 3.5 cm, centered at the fish head position. Particle acceleration in vertical direction and orthogonal to the direction of stimulation was neglected in this study.

Newton’s second law of motion, 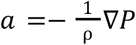 (pressure gradient force), links the spatial pressure gradient to particle acceleration. In water, with density ρ = 1,000 kg m−3 and speed of sound c = 1,500 m s−1, the following approximation holds for pressure signal frequencies f ≪ 100 kHz, where the pressure gradient is sampled at two locations |*x*_1_-*x*_2_| = 3.5 cm: 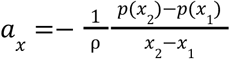. The approximation holds for all frequencies used in this experiment.

#### Sound targeting

To create the exact target waveforms for each stimulus with respect to sound pressure and particle acceleration at the position of the fish head, all presented sounds were calibrated using the empirically determined impulse responses of our playback system as detailed in ^57^. For each calibration, we documented and saved the final speaker waveforms and resulting acoustic signals at the fish’s position alongside all experimental data.

In this study we focus exclusively on acoustically evoked activity in response to sound pressure and therefore set up our calibration system to target zero particle acceleration, which effectively eliminates particle motion at the fish’s head position. Typical reduction factors are > 20 x.

### Acoustic playback hardware

Sounds delivered by the data acquisition card were amplified by a commercial Apart Champ 4 4-channel power amplifier and played back with small speakers (Ekulit LSF-27M/SC, 0,03W, 8 Ω) in custom-designed, waterproofed and 3D-printed casings.

#### Acoustic stimuli

All sound pressure stimuli were delivered at a peak-to-peak amplitude of 44 Pascal. The specific stimulus sets used in this study are described below.

The data presented in Figures 1 and 2 are based on a stimulus set containing pulse bursts and pure tones. The pulse bursts consisted of Gaussian double pulses at a center frequency of 5 kHz with < 1 ms duration, presented for 2 seconds at the following pulse repetition rates: 4, 8, 15, 30, 60, 120, 250, 500, and 1000 Hz. The pure tones were 2 seconds in duration with 20 ms of smooth onset and offset ramps, presented at the following quasi-logarithmically spaced frequencies: 150, 210, 300, 450, 610, 870, 1250, 1750, and 2500 Hz. Frequencies below 150 Hz were challenging to achieve due to the limited size and power range of the chosen speakers and were therefore excluded from this study.

The data presented in Figures 2d,e are based on a stimulus set that gradually morphs from a pulse burst (120 Hz repetition rate, Gaussian double pulses at 5 kHz center frequency) into a pure tone at 120 Hz, with each stimulus lasting 2 seconds. The stimulus waveforms were generated using the formula:

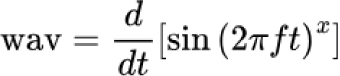

where *f* is the fundamental frequency (120 Hz), *t* is time, and *x* is an empirically determined exponent controlling the spectral shape. By varying the exponent x, we achieved spectral center frequencies (maxima of the power spectra) of 5000 (100%), 2644 (83%), 1440 (67%), 720 (48%), 360 (29%), 240 (19%), and 120 (0%) Hz. As x increases, the waveform transitions from a burst with prominent higher spectral content to a pure sinusoid.

The data presented in Figure 3 are based on a stimulus set containing 2-second pulse bursts (Gaussian double pulses at 5 kHz center frequency) with the following pulse repetition rates: 15, 30, 45, 60, 75, 90, 105, 120, 150, 200, and 250 Hz.

The data presented in Figures 4 are based on a stimulus set containing pulse bursts at 120 Hz repetition rate, where each burst contained the following pulse counts: 2, 4, 6, 8, 15, 30, 60, 120, 240, and 480.

### Analysis of imaging data

#### Segmentation and calculation of ΔF/F

We identified cell nuclei using local maxima detection applied to the local correlation map. Local correlation was calculated for each voxel relative to its 26-connected neighbors. This correlation map was then convolved with a 3D difference of Gaussians kernel (σ_0_ = 1 px, σ_1_ = 4 px). Local maxima were identified within spherical neighborhoods (⌀ 5 μm), approximately matching the size of cell nuclei. These detected maxima underwent global thresholding determined through visual inspection. Temporal traces for the remaining points were extracted by computing the mean of a Gaussian footprint (FWHM = 5 μm) positioned at each point. A baseline (running F_0_) for each trace was calculated through sequential filtering: median filter (7 s), followed by minimum filter (101 s moving window), and Gaussian filter (σ = 101 s). The ΔF/F was calculated relative to this baseline and then centered. Noise traces were rejected using a spectral SNR criterion exploiting the broad-band nature of shot noise versus the lower-frequency concentration of signals (we computed the ratio of average power below 0.25 Hz to average power at or above 0.25 Hz; traces were included if this ratio was ≥ 1.3). To eliminate artifacts from cells near the field of view edges that appeared intermittently due to fish motion, we excluded traces with dF/F values falling outside − 100%…2000%. All subsequent analyses used this processed data. For 2D activity maps (Fig. 1d), we used rastermap (v1.0; https://github.com/mouseland/rastermap; ^58^) with default parameters to display traces arranged by similarity.

#### Regression analysis

We used multivariate linear regression to quantify stimulus-evoked responses across neurons. For each stimulus and trial, we constructed a rectangular regressor based on its onset and offset frames, normalized to unit sum across frames. These regressors were convolved with an exponential kernel (τ = 4 s) to account for the empirically determined rise and decay kinetics of the calcium fluorescence signals. We assessed the quality of fit using the Pearson correlation coefficient *r* in two complementary ways. First, we correlated each neuron’s response trace with all individual trial regressors (n_stimuli × n_trials) to determine the overall goodness of fit (*r*_trials_). Second, we correlated response traces with a single regressor per stimulus (pooling across all trials) to quantify the overall reliability and entrainment of neuronal responses to each stimulus (*r*_stimuli_). To identify neurons with robust responses to acoustic stimulation, we applied selection criteria of *r*_trials_ > 0.7 and *r*_stimuli_ > 0.5, and used all neurons meeting these thresholds for subsequent analyses. Unless stated otherwise, the resulting regression coefficients 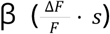 were used for all analyses. Because regressors are normalized to unit sum over time, β is directly proportional to the total exponential-kernel-weighted, integrated fluorescence response and has units of 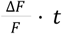. This ensures that response magnitude does not scale with stimulus duration, making coefficients comparable across stimuli of different lengths.

#### Best-frequency response curve analysis

We calculated region- and stimulus-specific response curves based on individual neurons’ best responses. For each neuron within a selected anatomical region, we identified the stimulus that elicited the strongest response. We then aggregated the response curves corresponding to each stimulus and computed the mean and standard deviation across neurons sharing the same preferred stimulus.

#### Calculation of pulse-tone-preference index

We quantified the relative response intensities to pulse bursts versus pure tones using a preference index ranging from −1 (exclusively burst-responsive) to 1 (exclusively tone-responsive). The preference index was calculated using the following formula: 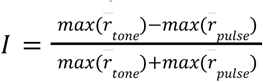, where max(r̄_tone_) is the maximum average response across all pure tone stimuli and max(r̄_pulse_) is the maximum average response across all pulse burst stimuli for each cell

#### Calculation of averaged correlation and covariance matrices

For each anatomically defined region, all neurons assigned to that region and their responses were selected. Pairwise similarity between stimuli was then quantified by computing the Pearson correlation coefficient between their corresponding population response vectors using scipy.stats.pearsonr (https://scipy.org). Repeating this procedure for all stimulus pairs yielded a stimulus–stimulus response similarity matrix for each anatomical region. The corresponding stimulus–stimulus covariance matrix was computed from the same population response vectors using numpy.cov (https://numpy.org).

#### Regional response magnitude distributions

For each anatomical region, neurons belonging to that region and their responses were selected based on ROI labels. For each stimulus, the distribution of response values across the selected neurons was summarized by computing specified percentiles using numpy.percentile. This was performed separately for each dataset, and the resulting stimulus-by-percentile matrices were subsequently averaged across datasets to obtain region-specific percentile response profiles.

#### Generation of replicate-averaged spatial response and effect-size maps

Neuronal responses were registered to a common 3D reference space for each recording and sex using neuron coordinates. Stimuli were grouped into short-burst and long-burst categories (short bursts: < 60 pulses, long bursts: > 60 pulses, where 60 pulses corresponds to a burst of 500 ms duration). For each category, voxelwise response volumes were generated by assigning neuronal responses to spatial locations in the shared sex-specific reference space and taking the maximum response across stimuli within the category, followed by Gaussian spatial smoothing (σ = 10 µm). For each sex, voxelwise mean and standard deviation maps were computed across recordings. Sex differences were quantified by calculating voxelwise standardized effect sizes between male and female response maps using the pooled standard deviation. For visualization, positive and negative effect sizes were separated into two channels and projected using a two-dimensional colormap (orange and blue), allowing simultaneous representation of male- and female-biased spatial effects.

#### Anatomical registration and cell mapping

To enable comparisons of neuronal activity patterns across recordings, sex, and stimuli, we registered each recording to sex-specific reference templates. These templates were generated in several steps. First, we computed an average volume for each recording by selecting and averaging the 100 best-correlating volumes. These recording-specific average brains were then registered to male and female OPM reference templates generated by pooling across many recordings ^21^. The OPM templates are registered to sex-matched anatomical two-photon templates, effectively bridging the functional data to the standardized reference brain space. Anatomical region labels obtained for each neuron were subsequently used for further analysis.

### Statistics

For the comparison of regional counts of responsive neurons (Fig. S11) we tested for significance differences between sexes using the Mann-Whithney U test. In our behavioral playback experiments statistical comparisons against sham and between sexes were made using Welch’s two-sample t-test.

### Data availability

Data underlying this study will be made available on gin.g-node.org upon publication.

### Competing interests

The authors declare no competing interests.

## Acknowledgements

We thank Constance Scharff, Livia de Hoz, Jan Benda, and Armin Bahl for helpful discussions and for feedback on this manuscript. We thank our fish facility team for excellent fish care and experimental support. We acknowledge support by the German Research Foundation (DFG, projects 432195732, 532521431 and EXC-2049-390688087), the European Research Council (ERC2021-CoG-101043615), the Einstein Foundation (EPP-2017-413), and the Alfried Krupp von Bohlen und Halbach Foundation.

## Supplemental figures

**Figure S1.**
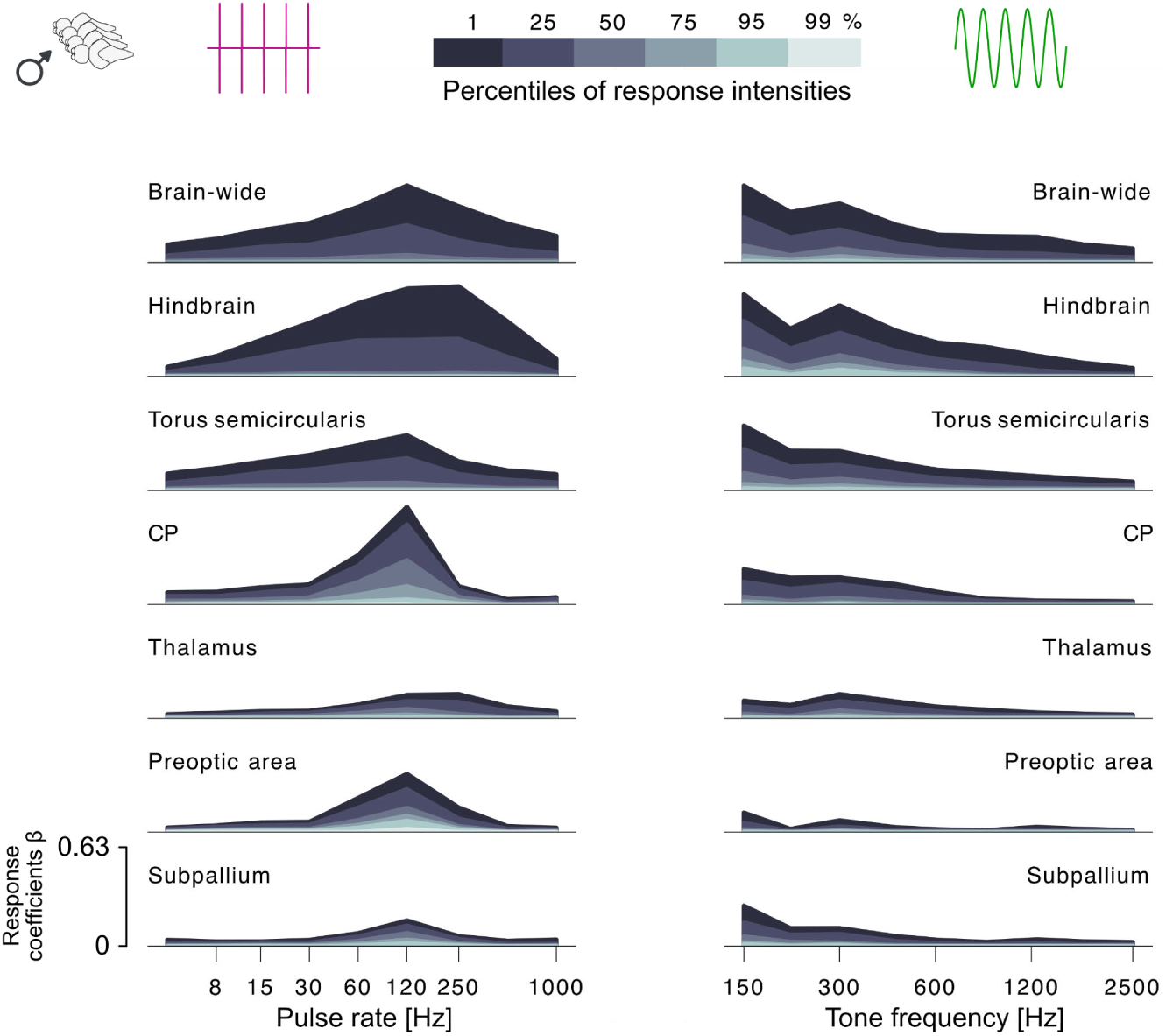
Brain-wide male sensitivity to pulsed and tonal sounds: Brain-wide, regional response magnitude distributions for pulsed (left) and tonal stimuli (right) in male fish. Percentile curves (1st–99th percentile) of stimulus-evoked response amplitudes by brain region, showing differential sensitivity across auditory regions. The Y-axis shows response coefficients 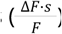.

**Figure S2.**
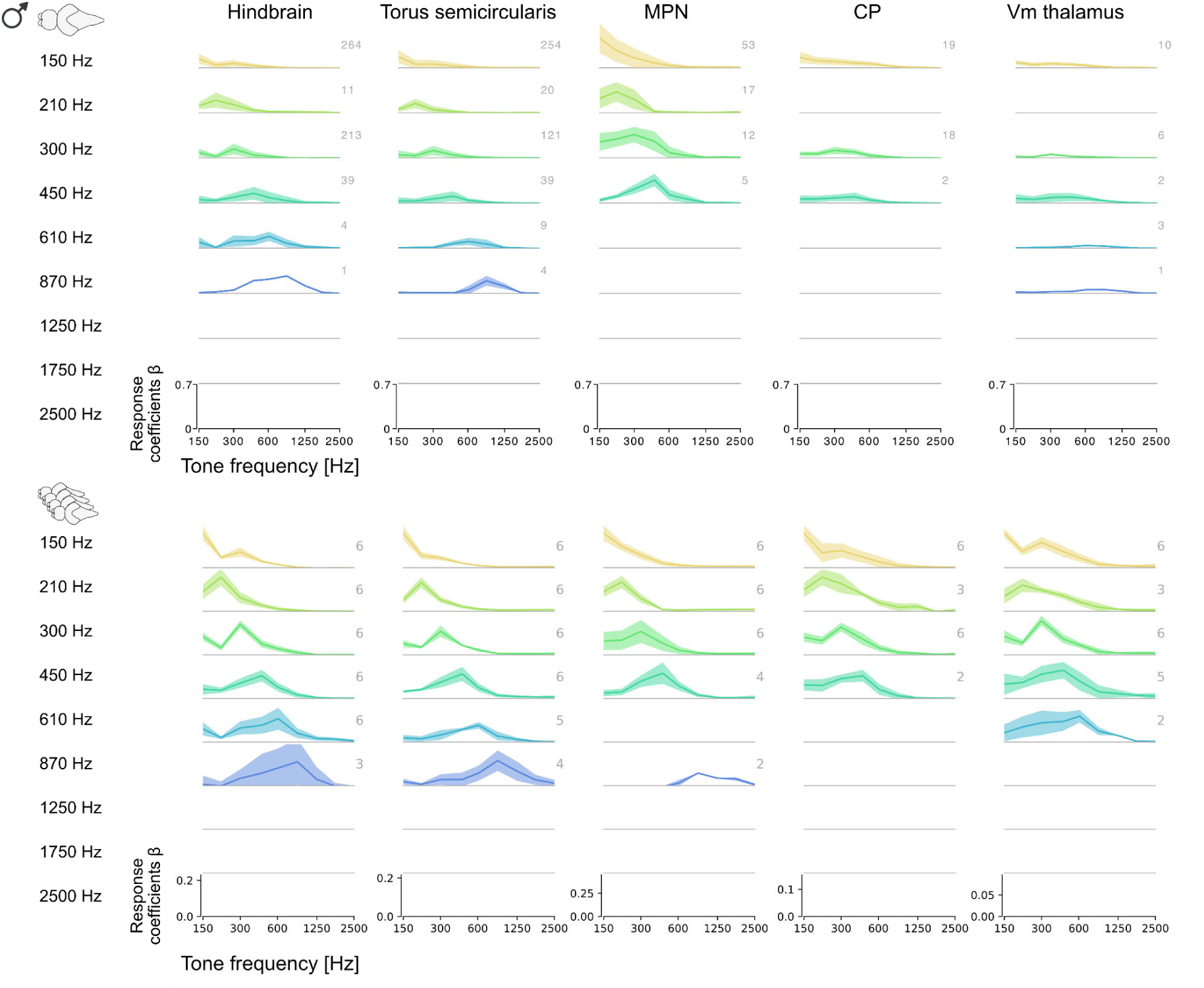
Male response curves to tonal sounds: Each panel shows the average best-response curves for each tone-frequency (rows) and region (columns). **a**, Data from a single example fish where each panel shows the mean and standard deviation (envelope) across neuronal best-response curves. Numbers next to the curves depict the number of cells in the aggregate curve. **b**, Aggregate across recordings (n=6), where each panel shows the mean and standard deviation across individual datasets. Numbers next to the curves depict the number of datasets in the aggregate curve. Datasets with less than two best-tuned cells per category were dismissed. The Y-axis shows response coefficients 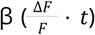.

**Figure S3.**
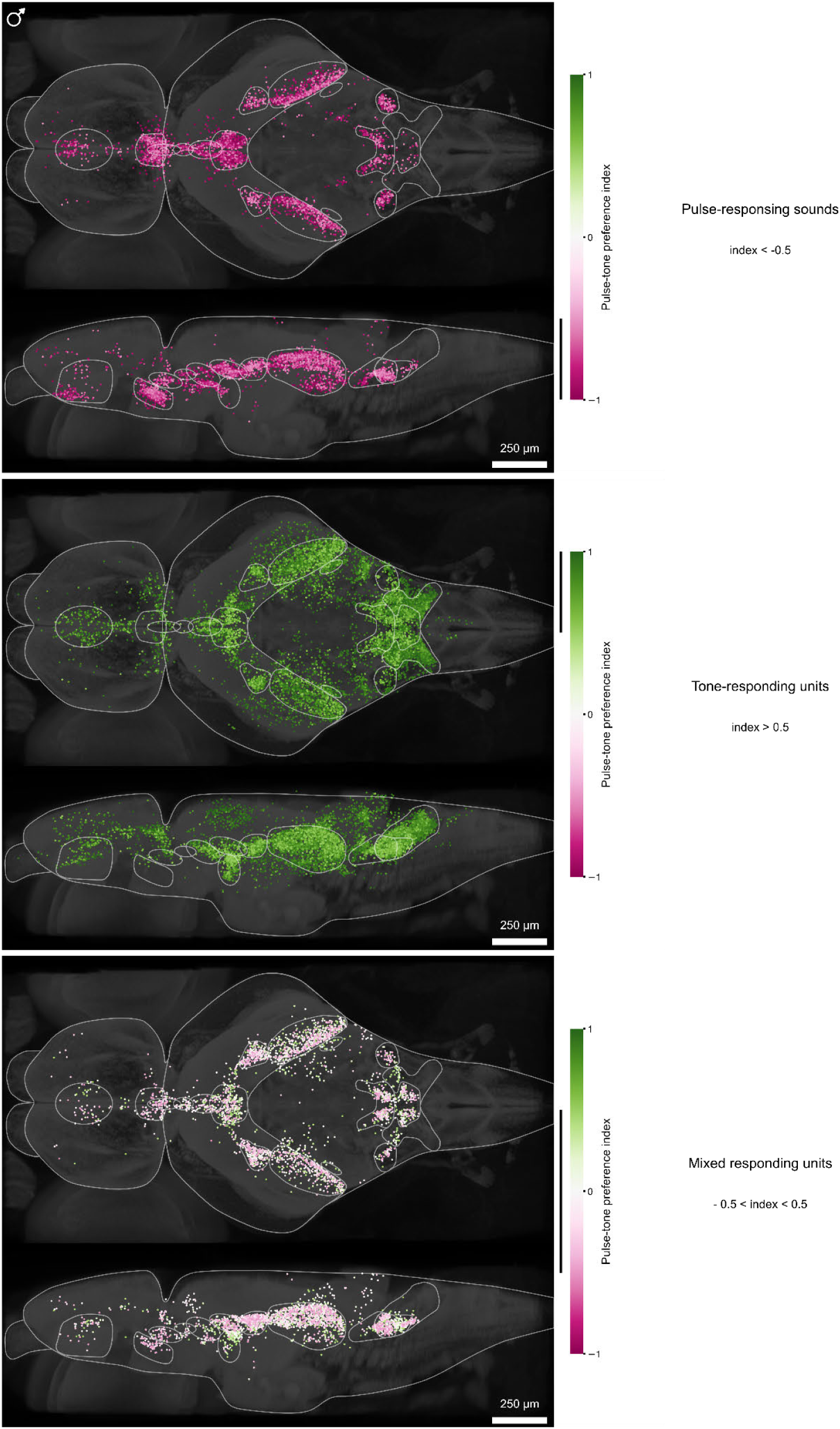
Male pulse / mixed / tone topography in detail: Spatial distribution of pulse-preferring (magenta) and tone-preferring (green) neurons aggregated across 6 fish and color-coded by pulse-tone preference index, shown separately for pulse-, tone-, and mixed selective units. The black bar next to the colorbar indicates the index-range shown in the respective plot.

**Figure S4.**
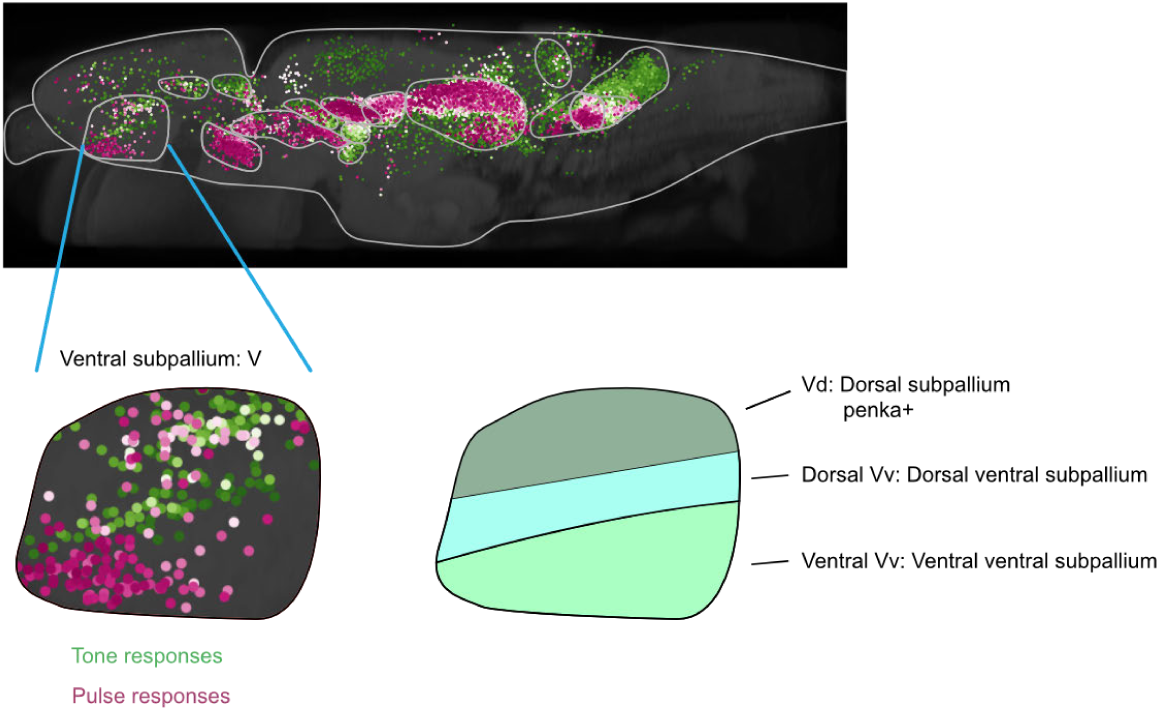
The subpallium exhibits functionally layered organization with sex-specific differences: The dorsal subpallium (Vd), homologous to the striatum ^48^, contains penka+ neurons ^48^ and shows mixed selectivity to both pulse and tone stimuli. The ventral subpallium (Vv), homologous in its entirety to the pallidum/septum ^48^, displays sexually dimorphic response patterns: In males, the Vv is functionally segregated – the dorsal Vv responds exclusively to pure tones, while the ventral Vv responds exclusively to long vocalization bursts. In females, the ventral subpallium does not respond to long vocalization bursts.

**Figure S5.**
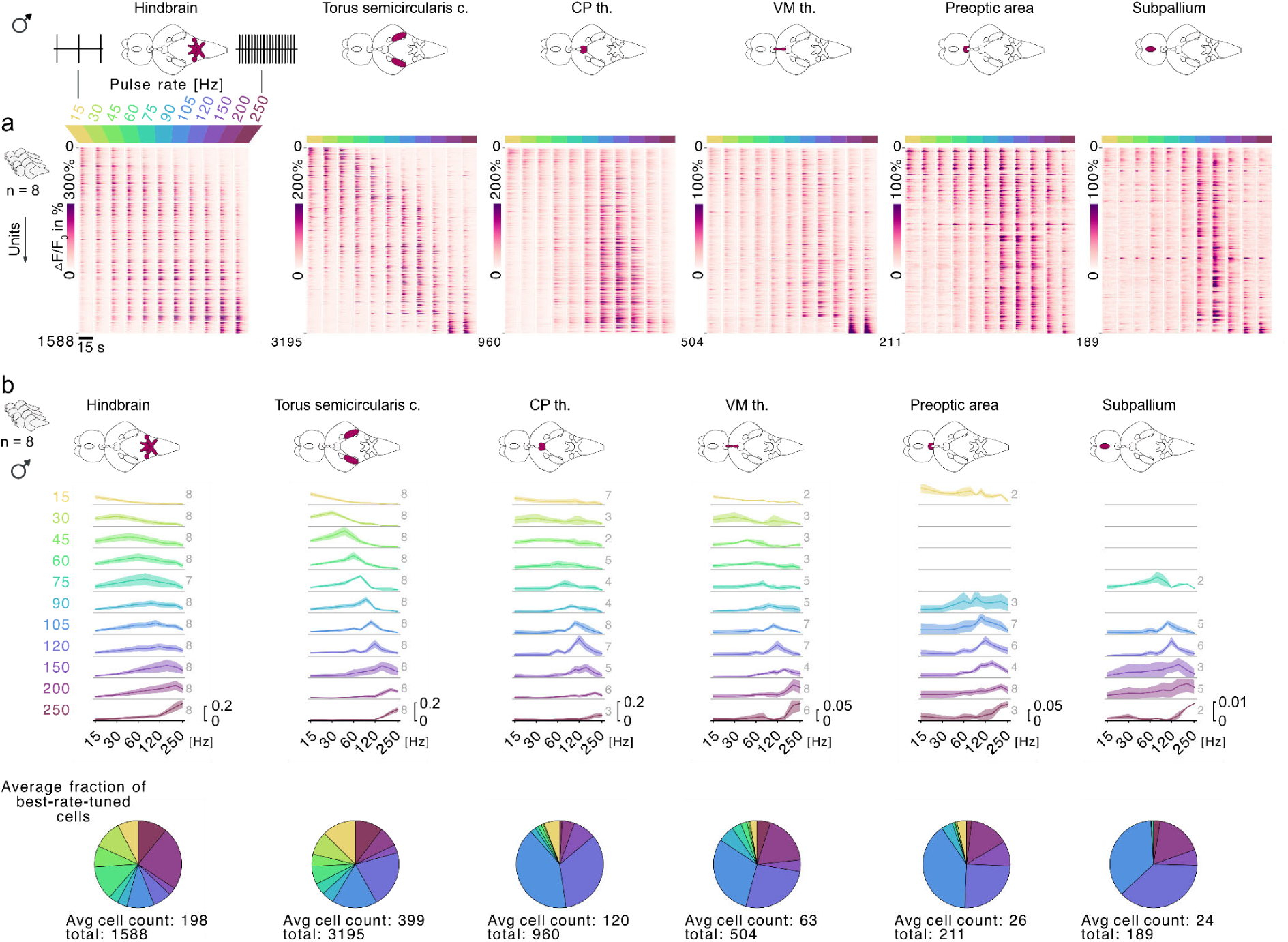
Male sensitivities to burst rate: **a**, Population-level regional response profiles. Heatmaps show trial-averaged calcium responses (ΔF/F) for individual neurons (rows) across pulse repetition rates from 15 to 250 Hz (columns) in six auditory brain regions (labeled above each panel), aggregated across 8 fish. Neurons are sorted by center-of-mass of individual response curves to reveal rate selectivity gradients. **b**, Male responses to burst rate aggregated across 8 recordings. Top: Each panel shows the average best-response curves for each burst rate (rows) and region (columns). Numbers next to response curves depict the number of datasets in the aggregate curve. Datasets with less than two best-tuned cells per category were dismissed. The Y-axis shows response coefficients 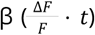. Bottom: Each pie chart shows the average fraction of best-tuned cells per stimulus per region.

**Figure S6.**
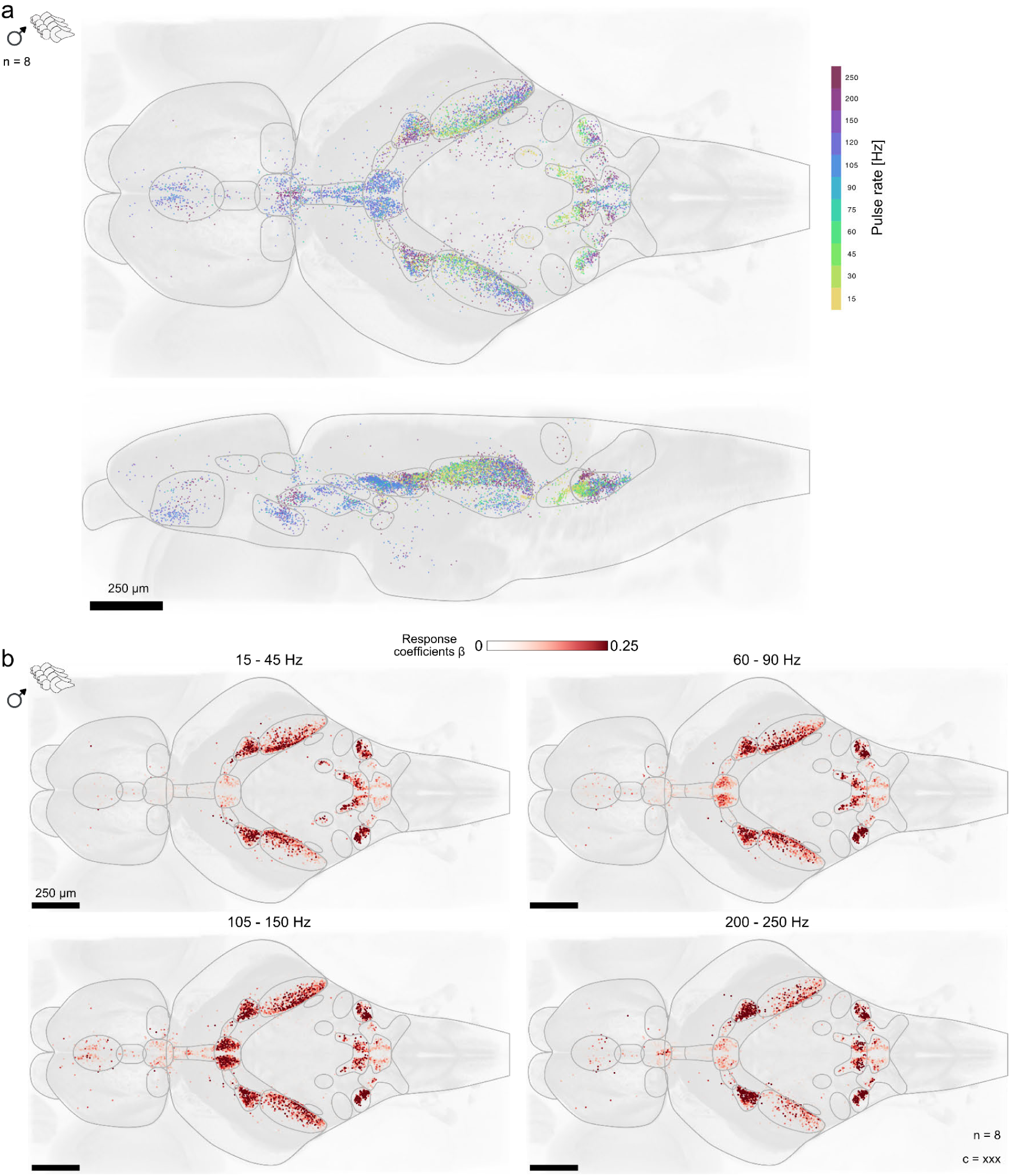
Male rate topology in detail: **a**, Orthogonal anatomical projections (top and side views) showing the spatial distribution of auditory neurons color-coded by their preferred pulse repetition rate (peak responses), revealing a topographic organization of repetition-rate selectivity. **b**, Top-view maps of auditory population responses (response coefficients 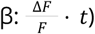) to four groups of pulse repetition rates. For each group, the map shows the maximum response evoked by any stimulus within that group.

**Figure S7.**
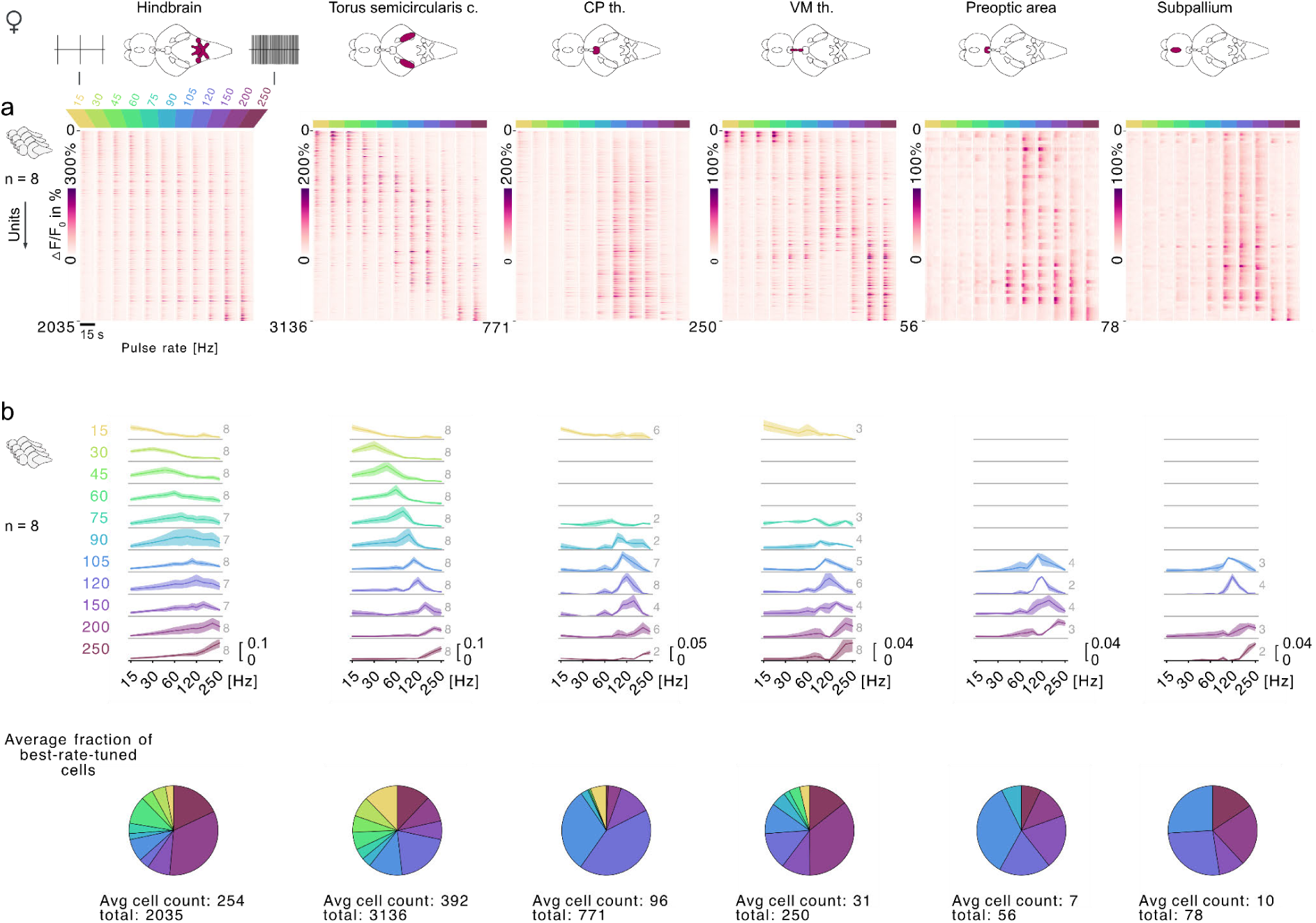
Female sensitivities to burst rate: **a**, Population-level regional response profiles. Heatmaps show trial-averaged calcium responses (ΔF/F) for individual neurons (rows) across pulse repetition rates from 15 to 250 Hz (columns) in six auditory brain regions (labeled above each panel), aggregated across 8 fish. Neurons are sorted by center-of-mass of individual response curves to reveal rate selectivity gradients. **b**, Female responses to burst rate aggregated across 8 recordings. Top: Each panel shows the average best-response curves for each burst rate (rows) and region (columns). Numbers next to response curves depict the number of datasets in the aggregate curve. Datasets with less than two best-tuned cells per category were dismissed. The Y-axis shows response coefficients 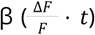. Bottom: Each pie chart shows the average fraction of best-tuned cells per stimulus per region.

**Figure S8.**
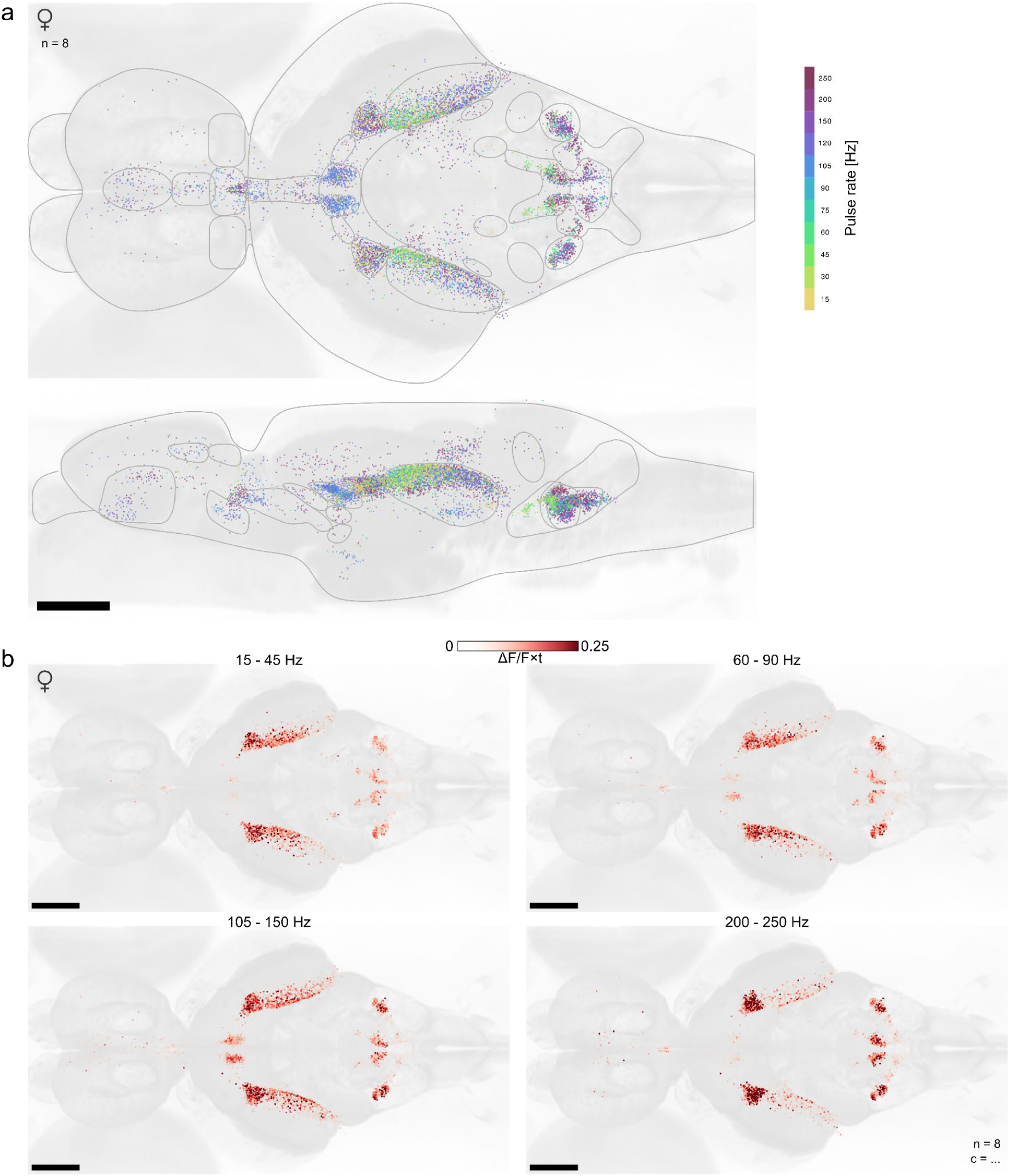
Female rate topology in detail: **a**, Orthogonal anatomical projections (top and side views) showing the spatial distribution of auditory neurons color-coded by their preferred pulse repetition rate (peak responses), revealing a topographic organization of repetition-rate selectivity. **b**, Top-view maps of auditory population responses (response coefficients 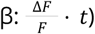 to four groups of pulse repetition rates. For each group, the map shows the maximum response evoked by any stimulus within that group.

**Figure S9.**
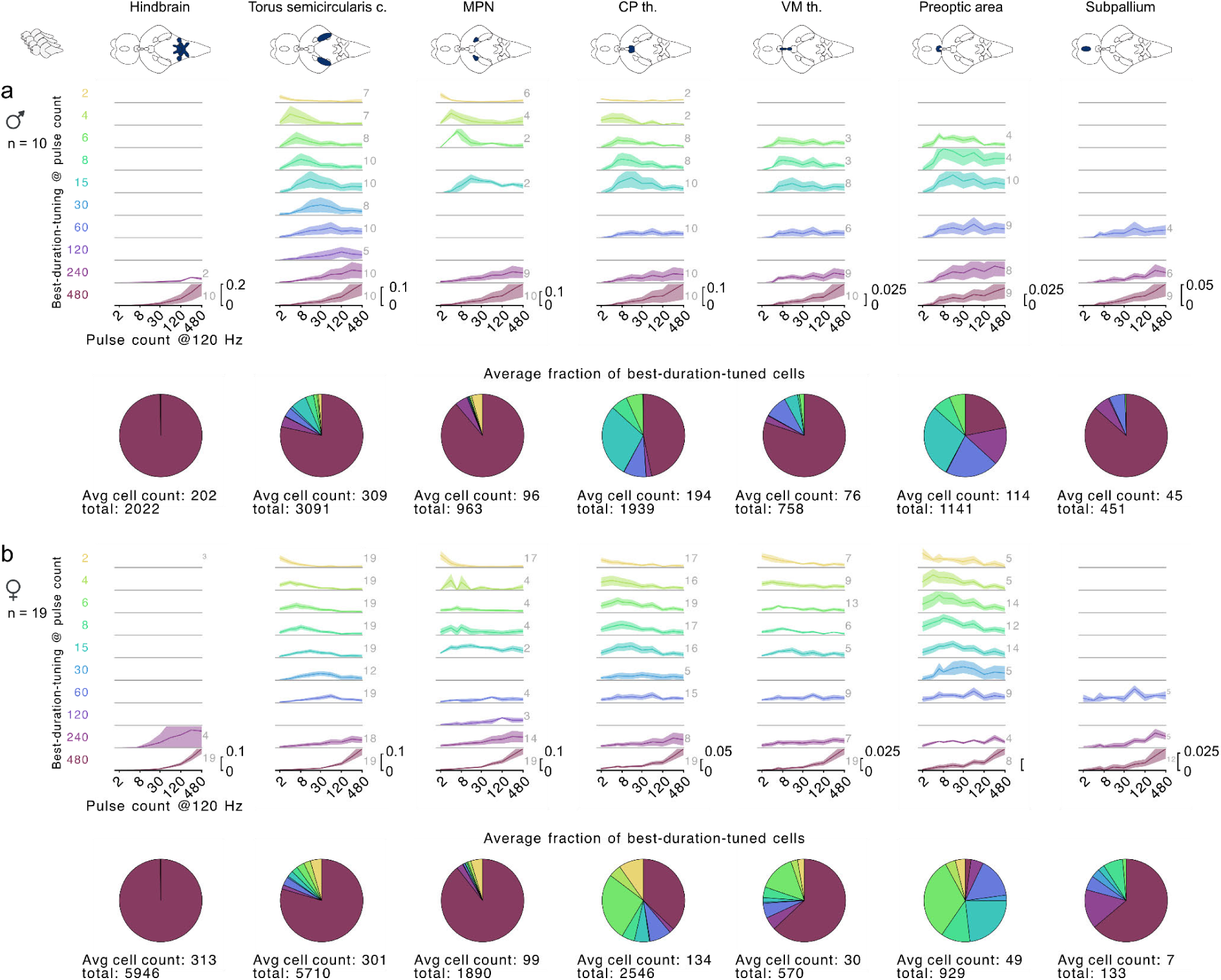
Male and female sensitivities to burst duration: **a**, Male responses to burst duration aggregated across 10 recordings. Top: Each panel shows the average best-responses curves for each burst duration (rows) and region (columns). Numbers next to the curves depict the number of datasets in the aggregate curve. Datasets with less than two best-tuned cells per category were dismissed. The Y-axis shows response coefficients 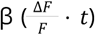. Bottom: Each pie chart shows the average fraction of best-tuned cells per stimulus per region. **b**, Female responses to burst duration aggregated across 19 recordings. Panel structure as in a).

**Figure S10.**
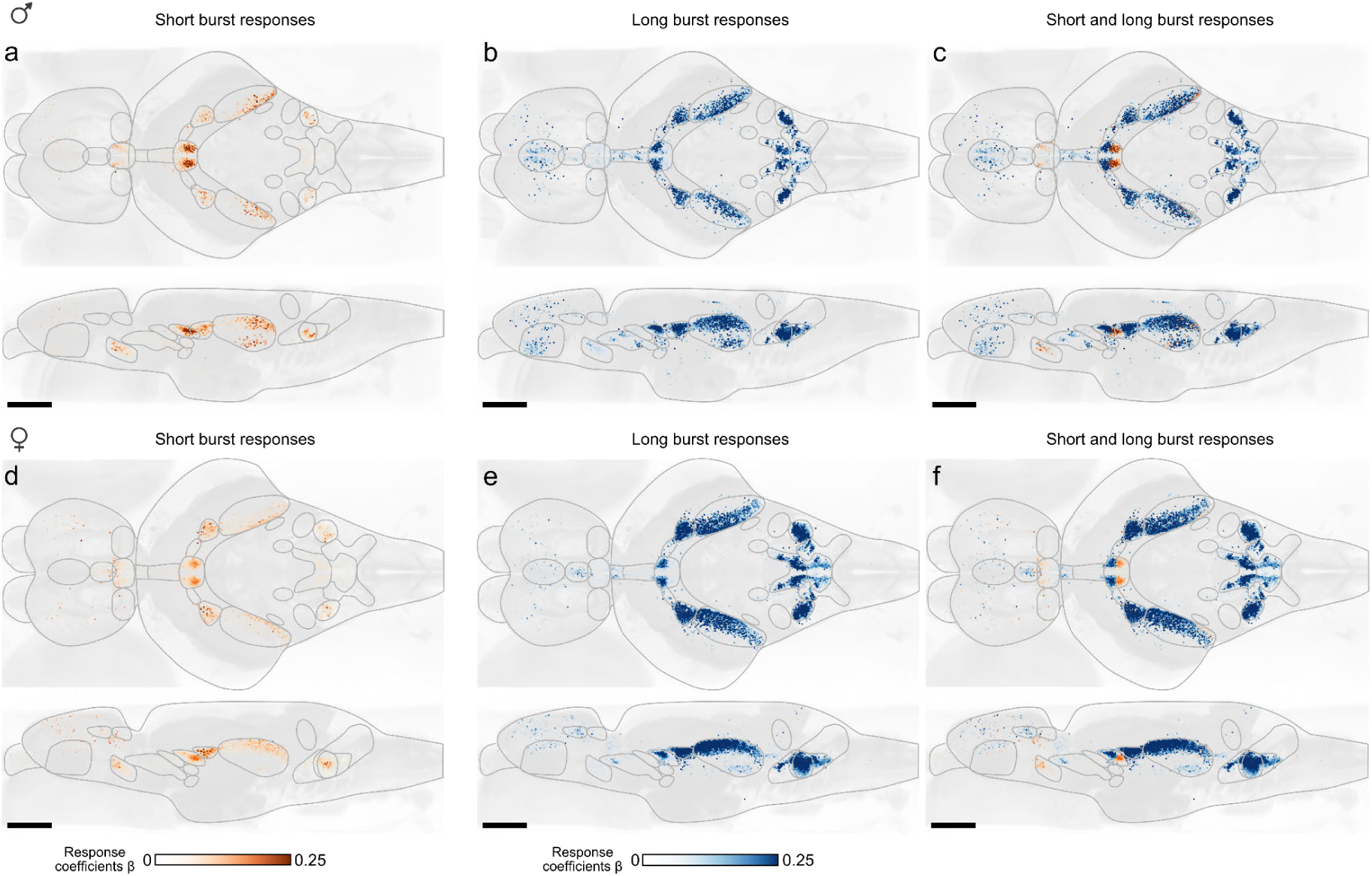
Males and females pooled responses to short and long duration bursts: **a,d**, Male and female responses to short bursts, pooled across recordings. **b,e**, Male and female responses to long bursts, pooled across recordings. **c,f**, Overlay of male and female responses to short and long bursts, pooled across recordings. Males, n=10; females, n=19.

**Figure S11.**
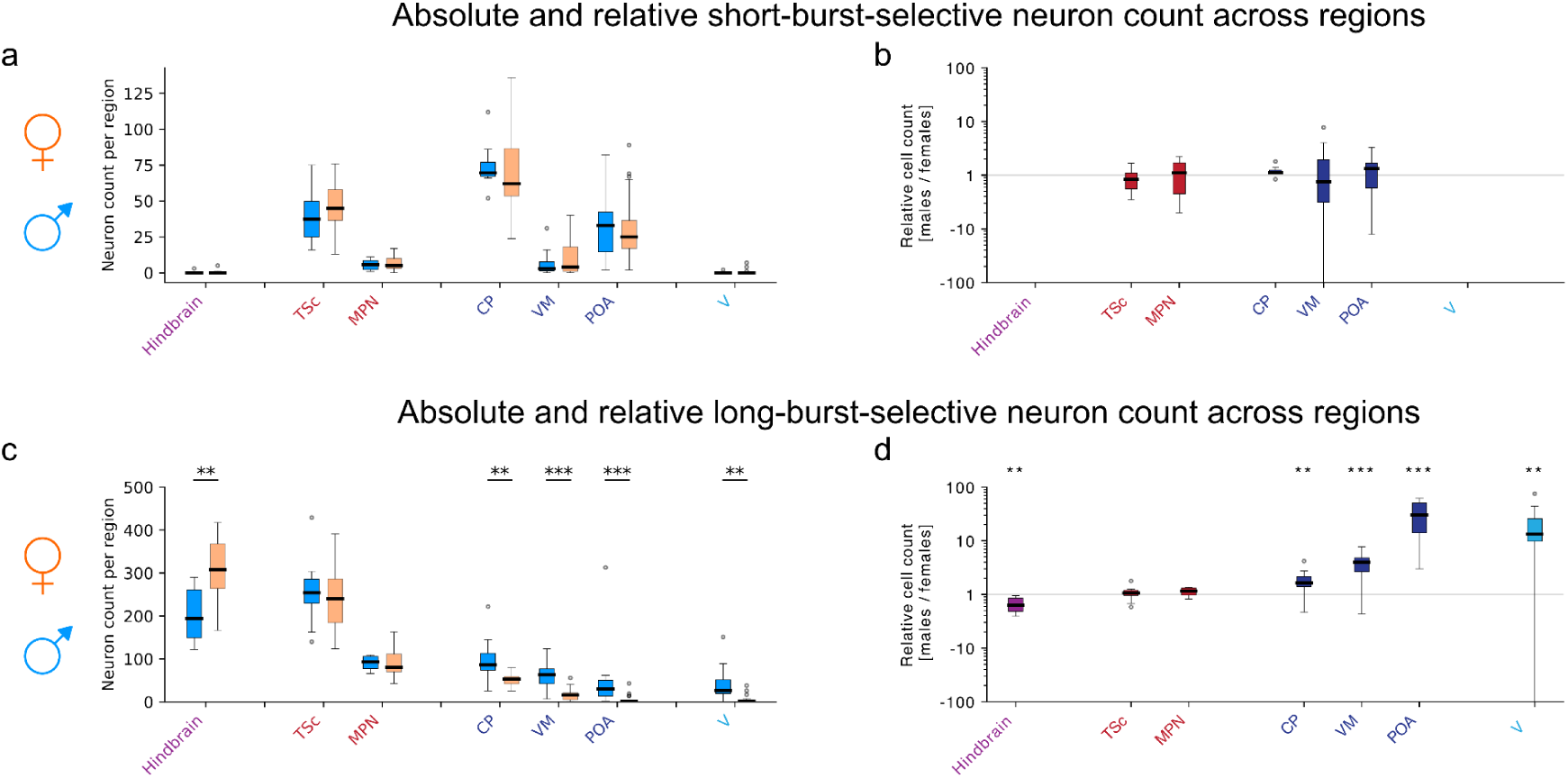
Sex differences in burst-duration-selective neuron distribution across brain regions: Regional distribution of neurons responsive to pulse bursts and selective for burst duration in males (n=10) versus females (n=19). **a**, Absolute counts of short burst-selective neurons. **b**, Relative short burst-selective neuron counts per region (male/female median ratio); values >1 indicate male bias, <1 indicate female bias, and ∼1 indicate similar distributions. **c**, Absolute counts of long burst-selective neurons. Regional counts were compared between sexes using a Mann-Whithney U test; significant differences were observed in the hindbrain, CP th., VM th., POA, and V (p=0.0016, 0.0016, 0.0009, 0.0002, and 0.0042, respectively). **d**, Relative long burst-selective neuron counts per region (male/female median ratio). Significance indicators refer to the same tests performed in **c**.

## Notes

### Competing Interest Statement

The authors have declared no competing interest.

### Summary of Updates

Addition of morph stimulus data (Figure 2). 7 new & 5 updated main-text panels.

